# p75NTR prevents the onset of cerebellar granule cell migration via RhoA activation

**DOI:** 10.1101/2022.04.26.489604

**Authors:** Juan P. Zanin, Wilma J. Friedman

**Author notes:** Corresponding author, Juan Zanin.

## Abstract

Neuronal migration is one of the fundamental processes during brain development. Several neurodevelopmental disorders can be traced back to dysregulated migration. Although substantial efforts have been placed in identifying molecular signals that stimulate migration, little is known about potential mechanisms that restrict migration. These restrictive mechanisms are essential for proper development since it helps coordinate the timing for each neuronal population to arrive and establish proper connections. Moreover, preventing migration away from a proliferative niche is necessary to maintain a pool of proliferating cells until the proper number of neuronal progenitors is attained.

Here, we identify an anti-migratory role for the p75 neurotrophin receptor (p75NTR) in cerebellar development. Our results show that granule cell precursors robustly express p75NTR in the external granule layer (EGL) when they are proliferating during postnatal development, however, they do not express p75NTR when they migrate either from the rhombic lip during embryonic development or from the EGL during postnatal development. We show that p75NTR prevented GCP migration by maintaining elevated levels of active RhoA. The expression of p75NTR was sufficient to prevent the migration of the granule cells even in the presence of BDNF, a well-established chemotactic signal for this cell population. Our findings suggest that the expression of p75NTR might be a critical signal that stops and maintains the GCPs in the proliferative niche of the EGL, promoting the clonal expansion of cerebellar granule neurons.

## Introduction

Neuronal migration is a fundamental process in nervous system development. Several neurological disorders can be traced back to defects in migration during development (Buchsbaum & Cappello, 2019). In mammals, a highly coordinated series of neuronal migration events are essential for the establishment of the layered structure of the brain, while bringing neurons into the correct location to assemble the proper neuronal connections.

The cerebellum is well-known for its role in motor control, yet it is also involved in non-motor functions such as decision making (Deverett et al., 2018, 2019), reward anticipation (Kim et al., 2017), and social interaction (Carta et al., 2019). Cerebellar granule cells are a unique neuronal population since they undergo two rounds of proliferation/migration cycle during development. In the first round, starting at E10.5 in rats, cells proliferate in the rhombic lip, and around E12.5 the granule cell precursors (GCPs) begin to migrate tangentially on the surface of the future cerebellum, where they will stop migrating and establish a transient proliferative zone, the external granule layer (EGL)(Altman & Bayer, 1996). In the EGL, the GCPs start the second round of proliferation, undergoing clonal expansion during the first 2-3 postnatal weeks. During this entire period, waves of GCPs exit the cell cycle and start the second migratory stage. Using the Bergman glia as a track together with molecular cues, the GCPs pass through the Purkinje cell layer (PC) and colonize the internal granule layer (IGL), their final destination. Although several studies have identified signals that stimulate and guide granule cell migration, little is known about potential mechanisms that prevent the onset of migration. These restrictive mechanisms are critical for proper development since they help to coordinate the proper timing of cell migration away from a proliferative niche such as the EGL (Altman & Bayer, 1996).

The p75 neurotrophin receptor (p75NTR) can mediate a broad array of functions depending on the cellular context. During cerebellar development, p75NTR is highly expressed in the proliferating GCPs located in the outer EGL (oEGL) and is downregulated before the GCPs start migrating (Carter et al., 2003; Zanin et al., 2016). Among the multiple signaling proteins that are regulated by p75NTR, RhoA regulation is particularly interesting, since p75NTR can maintain elevated levels of active RhoA in a ligand-independent manner (Yamashita et al., 1999; Yamashita & Tohyama, 2003). RhoA is a member of the small sub-family of Rho-GTPases that cycle between a GTP-bound (active) and a GDP-bound (inactive) form. Through this mechanism, these molecules can coordinate multiple aspects of cellular responses. In particular, due to their regulation of cytoskeletal dynamics, these GTPases are involved in different aspects of neuronal migration (Xu et al., 2019). Therefore, disruption of Rho-GTPase activity has been associated with migration defects leading to several behavioral and developmental disorders (Boettner & Van Aelst, 2002; Govek et al., 2005). Regulation of this signaling pathway suggests the possibility that p75NTR can influence neuronal migration via the regulation of RhoA activity.

In the present study, we aimed to investigate the role of p75NTR in regulating cerebellar granule cell (CGN) migration. We found that this receptor exerts an anti-migratory effect on the granule cells, both during embryonic and postnatal development, and must be downregulated to allow the CGNs to migrate away from the EGL. Ex-vivo analysis showed that overexpression of either p75NTR or activated RhoA blocks the migration of granule cells. Our findings reveal a novel role for p75NTR in preventing neuronal migration by maintaining RhoA activation in the developing cerebellum.

## Results

### P75NTR is expressed in mitotic, non-migrating GCPs

During early embryonic development (E13.5) the precursors of the granule cells, indicated by the expression of PAX6 (Fig 1 A-B and Fig S1 green), begin to disperse rostrally from the rhombic lip, interestingly these precursor cells do not express the neuronal migratory marker DCX, suggesting these cells did not yet acquire a neuronal fate (Fig 1 A-B and Fig S1 red). During this period, no expression of p75NTR was observed in the migrating cells (Fig 1 A-B and Fig S1 white). At later stages (E15.5) the majority of the precursor cells still migrate tangentially to complete the formation of the EGL. However, a subpopulation of cells begins to emerge, located in the internal part of the migratory stream, these cells express PAX6 (Fig 1 A-B and Fig S1 green) and the neurotrophin receptor p75NTR (Fig 1 A-B and Fig S1 white). At the final stages of embryonic development (E17.5), there is an enlargement of the EGL, which is accompanied by an increase in the number of p75NTR expressing cells (Fig 1A-B and Fig S1 white), which continues throughout postnatal development until the EGL disappears between 2 and 3 weeks after birth.

**Figure 1:**
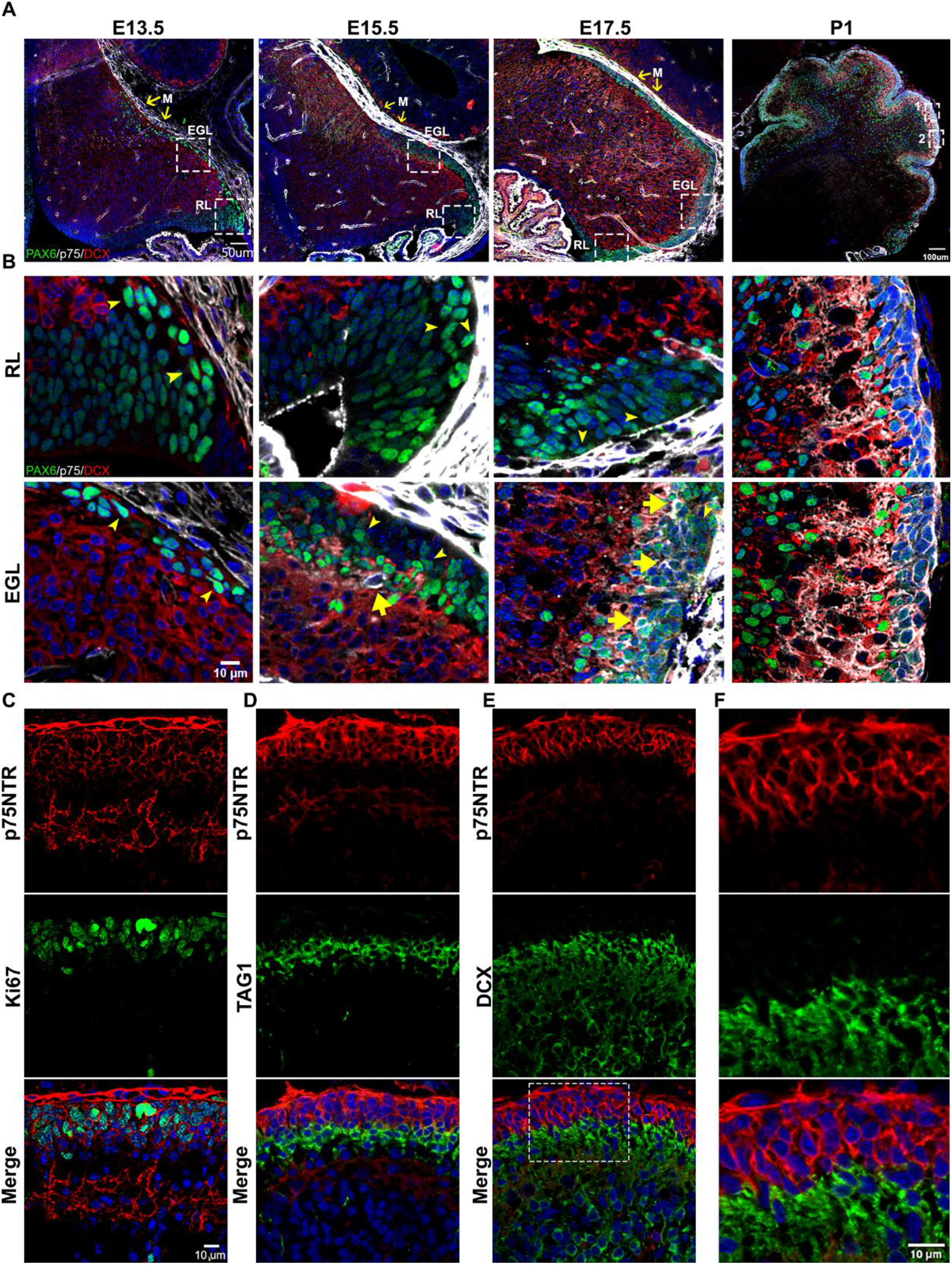
Developmental expression of p75NTR in the cerebellum. (A) Developmental expression of p75NTR (white), Ki67 (red), and Pax6 (green) at the indicated ages. Note the high level of p75NTR in the meninges as well as the developing granule cell progenitors. Yellow arrows indicate the meninges – M, RL-rhombic lip, EGL – external granule layer. (B) High magnification of the inset showed in A. Cells expressing p75NTR (arrows), migrating cells negative for p75NTR (arrowheads). (C) Expression of p75NTR (red) and Ki67 green in the cerebellum of P7 mouse pups. (D) Immunohistochemistry of the expression of p75NTR (red) and TAG1 (green) in the cerebellum of P7 mouse pups. (E) Immunohistochemistry of the expression of p75NTR (red) and DCX (green) in the cerebellum of P7 mouse pups. (F) High magnification of the inset showed in E.

During postnatal development, p75NTR is expressed in the GCPs throughout the entire EGL (Fig 1C-F, Red). Moreover, the expression of p75NTR closely correlates with the proliferation levels in the EGL, with a peak of p75NTR expression during the maximal proliferation stage of the GCPs, which in mice occurs around P5-7 (Fig. 1C green). Postmitotic GCPs, located mostly in the internal part of the EGL, begin to express neuronal differentiation and migratory markers such as TAG1 (Fig. 1D, Green) or DCX (Fig 1E, Green) in preparation for the radial migratory phase towards the internal granule layer (IGL). In contrast to the expression of p75NTR in proliferating cells, postmitotic cells downregulate the expression of p75NTR, and no overlap between p75NTR and the migratory markers was observed (Fig. 1D-F). This well-defined boundary between the proliferating cells expressing p75NTR and the migrating TAG1 or DCX positive cells indicates that exiting the cell cycle is accompanied by a reduction in p75NTR expression in GCPs as they leave the cell cycle. The negative correlation between the cells that express p75NTR and the migratory cells observed at embryonic as well as postnatal stages suggests that the expression of the receptor could be a signal that stops the migration of cells coming from the rhombic lip, and retains the cells in the EGL until they are ready to migrate radially towards the IGL

### Shh induces GCP proliferation and maintains a high expression of p75NTR

Although there is a clear transition between the proliferating cells expressing p75NTR and the differentiating cells that downregulate the expression of the receptor, the specific function of the receptor in this transition is not clear. For instance, it is unclear whether p75NTR is required to keep the GCPs in the cell cycle, or whether the cells maintain the expression of p75NTR because they are in the cell cycle. To address this question, we took advantage of the spontaneous differentiation that the GCPs undergo in the absence of a mitogen in culture. In the presence of Shh, a well-known mitogen for GCPs (Kenney & Rowitch, 2000; Wallace, 1999), these cells are maintained in a proliferative and undifferentiated state with high expression levels of the proliferation marker PCNA (Fig 2A, B). However, as early as 24 h in culture without Shh there is a significant reduction in the level of PCNA. We also observed that p75NTR expression significantly decreased in conditions without Shh, but remained elevated in the proliferative environment induced by Shh (Fig 2A, C). To further explore this positive correlation between proliferation levels and p75NTR expression, we exposed GCP cultures to different concentrations of Shh for 48 h (Fig 2D-K). Shh induced a dose-dependent increase in proliferation, estimated by the number of Ki67+ cells (Fig 2G top panels, 2H) and the relative increase in PCNA expression (Fig 2D, E). At the same time, Shh maintained high expression levels of p75NTR in a dose-dependent manner (Fig 2G middle panels, 2D, F, J). Consistent with the observations in the EGL during cerebellar development (Fig 1), the number of double-labeled cells expressing p75 and Ki67 also increased in response to Shh in a dose-dependent manner (Fig 2J). Furthermore, all the proliferating cells also expressed p75NTR, even in the control conditions where the few remaining proliferating cells also expressed p75NTR (Fig 2K).

**Figure 2:**
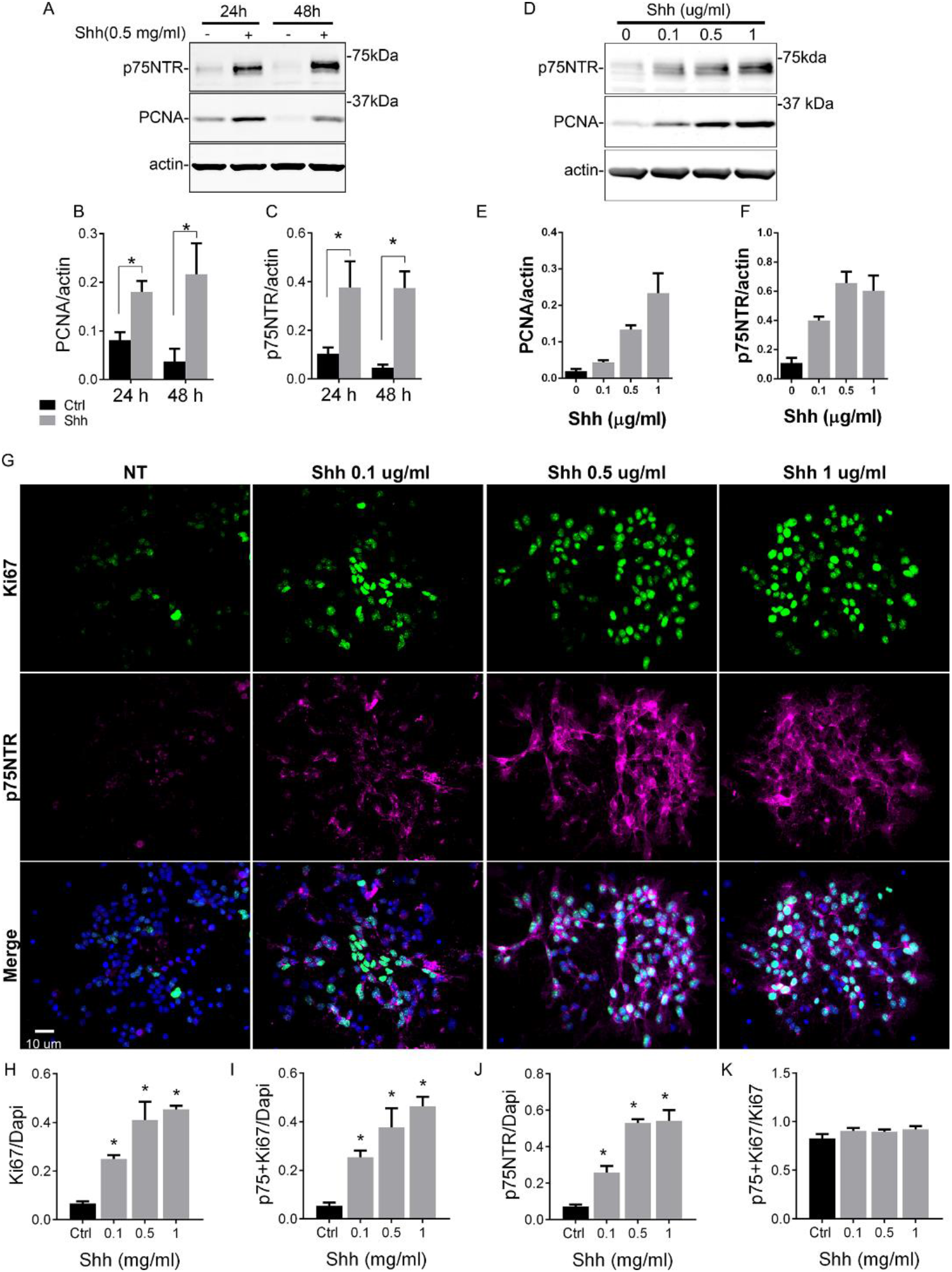
Shh regulates CGN proliferation and the expression of p75NTR. (A) Western Blot analysis of the temporal expression of p75NTR and PCNA (proliferation marker) in granule cell cultures exposed to Shh. (B and C) Quantification of PCNA and p75NTR expression relative to actin. Two-way ANOVA, N = 3, error bars indicate SEM. (D) Western Blot analysis of the expression of p75NTR and PCNA in granule cell cultures in response to increasing concentrations of Shh for 48h. (E) and F) Quantification of PCNA and p75NTR expression relative to actin. One-way ANOVA, N = 3, error bars indicate SEM. (G) Immunocytochemistry analysis of the expression of p75NTR and Ki67 in granule cell cultures in response to increasing concentrations of Shh for 48h. (H) Quantification of the total number of proliferating cells expressed as the percentage of cells expressing Ki67 over the total number of cells (Dapi). One-way ANOVA, N = 3, error bars indicate SEM. (I) Quantification of the number of p75NTR positive cells, expressed as the number of p75NTR positive cells over the total number of cells (Dapi). One-way ANOVA, N = 3, error bars indicate SEM. (J) Quantification of the number of Ki67+p75NTR double-label cells express as the number of cells expressing Ki67 and p75NTR over the total number of cells (Dapi). One-way ANOVA, N = 3, error bars indicate SEM. (K) Quantification of the number of proliferating cells that also express p75NTR, expressed as the number of double-label Ki67+p75NTR cells over the total number of cells expressing Ki67. One-way ANOVA, N = 3, error bars indicate SEM.

### Cell cycle exit correlates with a reduction in p75NTR levels

Previously, we have shown that proNT-3 can induce cell cycle exit of GCPs via p75NTR, even in the presence of Shh (Zanin et al., 2016). To determine whether inducing cell cycle exit of GCPs also elicits downregulation of p75NTR, we treated the cells with proNT-3 in the presence of the Shh analog SAG. Confirming the cell cycle withdrawal induced by proNT-3, we observed a reduction in the Ki67+ cells, which was accompanied by a significant decrease in p75NTR expression (Fig 3A-C). Consistent with increased differentiation, proNT-3 induced an increase in the levels of βIII-tubulin even in the presence of SAG (Fig 3A and 3D).

**Figure 3:**
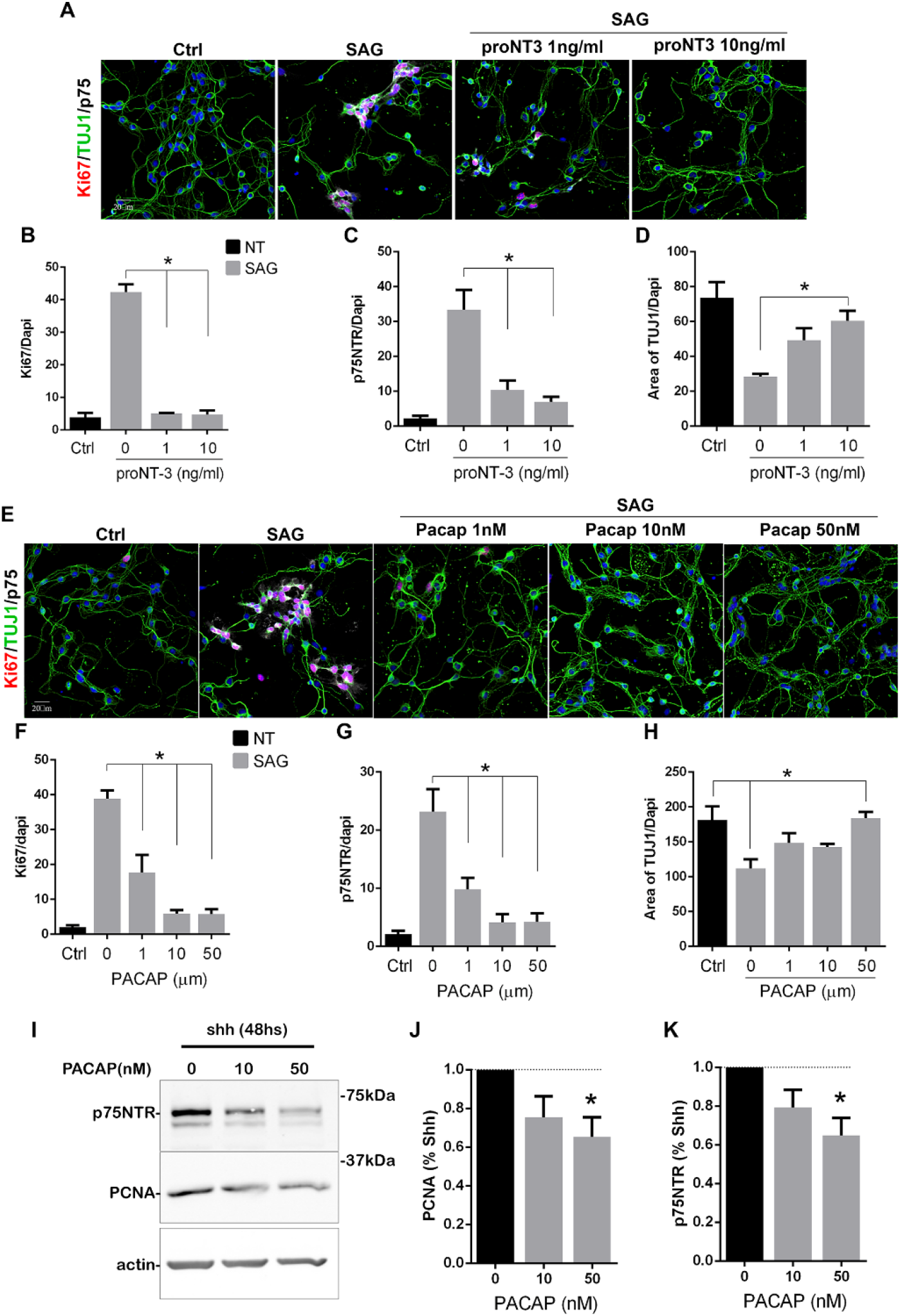
Cell cycle exit induced p75NTR downregulation. (A) Immunocytochemistry analysis of the expression of p75NTR (white), Ki67 (red), and βIII tubulin (green) in granule cell cultures in response to increasing concentrations of proNT-3 after 48h in culture. (B and C) Quantification of the number of cells expressing p75NTR or Ki67 over the total number of cells (Dapi). One-way ANOVA, N = 4, error bars indicate SEM. (D) Quantification of cerebellar granule cell differentiation expressed as the total area of processes positive for βIII tubulin over the total number of cells (Dapi). One-way ANOVA, N = 4, error bars indicate SEM. (E) Immunocytochemistry analysis of the expression of p75NTR (white), Ki67 (red), and DCX (green) in granule cell cultures in response to increasing concentrations of PACAP after 48h in culture. (F and G) Percentage of cells expressing Ki67 or p75NTR over the total number of cells (Dapi). One-way ANOVA, N = 4, error bars indicate SEM. (H) Quantification of cerebellar granule cell differentiation expressed as the total area of processes positive for DCX over the total number of cells (Dapi). One-way ANOVA, N = 4, error bars indicate SEM. (I) Western blot analysis of the expression of p75NTR and PCNA in granule cell cultures in response to increasing concentrations of PACAP after 48h in culture. (J and K) Quantification of PCNA or p75NTR expression in CGN cultures expose to PACAP and normalize to Shh alone. One-way ANOVA, N = 4, error bars indicate SEM.

To determine whether other signals that promote cell cycle withdrawal of GCPs also trigger a decrease in p75NTR expression, we treated granule cells with PACAP, a known anti-mitogenic ligand for GCPs (Nicot et al., 2002). We cultured GCPs for 48 h in the presence of SAG with and without PACAP and quantified their proliferation either by Ki67 staining or PCNA levels using Western blots (Fig 3E-K). PACAP induced a dose-dependent reduction in Ki67 (Fig 3E, F) and PCNA levels (Fig 3I-J) and a concomitant reduction in p75NTR expression in a dose-dependent manner, even in the presence of SAG (Fig 3E, G, I, and K). The reduced proliferation induced by PACAP was accompanied by increased neuronal differentiation, indicated by an increase in the levels of βIII-tubulin (Fig 3E, H).

### P75NTR overexpression does not promote cell cycle re-entry

Our data have demonstrated a positive correlation between the expression of p75NTR and the proliferating GCPs, and a complete absence of the receptor upon cell cycle exit. To directly test whether the expression of p75NTR is sufficient to maintain the GCPs in a state permissive for proliferation even after withdrawal of Shh, which would allow the cells to re-enter the cell cycle upon re-addition of Shh, we transfected GCPs with a p75-GFP construct to maintain expression of the receptor even after Shh was withdrawn (Figure 4). GCPs were then cultured in the absence of Shh for 24h, which is sufficient to reduce the levels of proliferation and the endogenous expression of p75NTR (see Fig. 2). The Shh analog SAG was then added for 24h and the proliferation levels were evaluated after the total 48h in culture. If p75NTR expression was sufficient to maintain the cells in a permissive proliferative state, we would expect to see an increase in proliferation with the re-exposure to Shh. As expected, cells that were maintained continuously in the presence of SAG for 48h continued to proliferate, regardless of the transfected construct (i.e. Ctrl-GFP or p75-GFP. Fig 4A right panels, 4B-F) similar to the response observed in nontransfected cells (Fig 2 and 4D-F). Similarly, both transfected and non-transfected cells cultured in the absence of the mitogen for 48 h did not proliferate (Fig 4A left panels, 4B-F). Interestingly, when cells were cultured without SAG for the first 24 h, and then with SAG for the next 24 h, there was no significant difference in the proliferation response between the Ctrl-GFP or p75-GFP transfected cells. Moreover, there was no difference between this condition and the cells maintained in the absence of SAG for the entire 48h (Fig 4A middle panels, 4B-F). Consistently, there was a significant reduction in proliferation levels in the cells maintained without the mitogen during the first 24h in culture, and the re-exposure to SAG did not induce re-entry to the cell cycle (Fig 4B, C, F), indicating that the continued expression of p75NTR is not sufficient to maintain the CGPs in a proliferative state, thus unresponsive to the delayed SAG addition.

**Figure 4:**
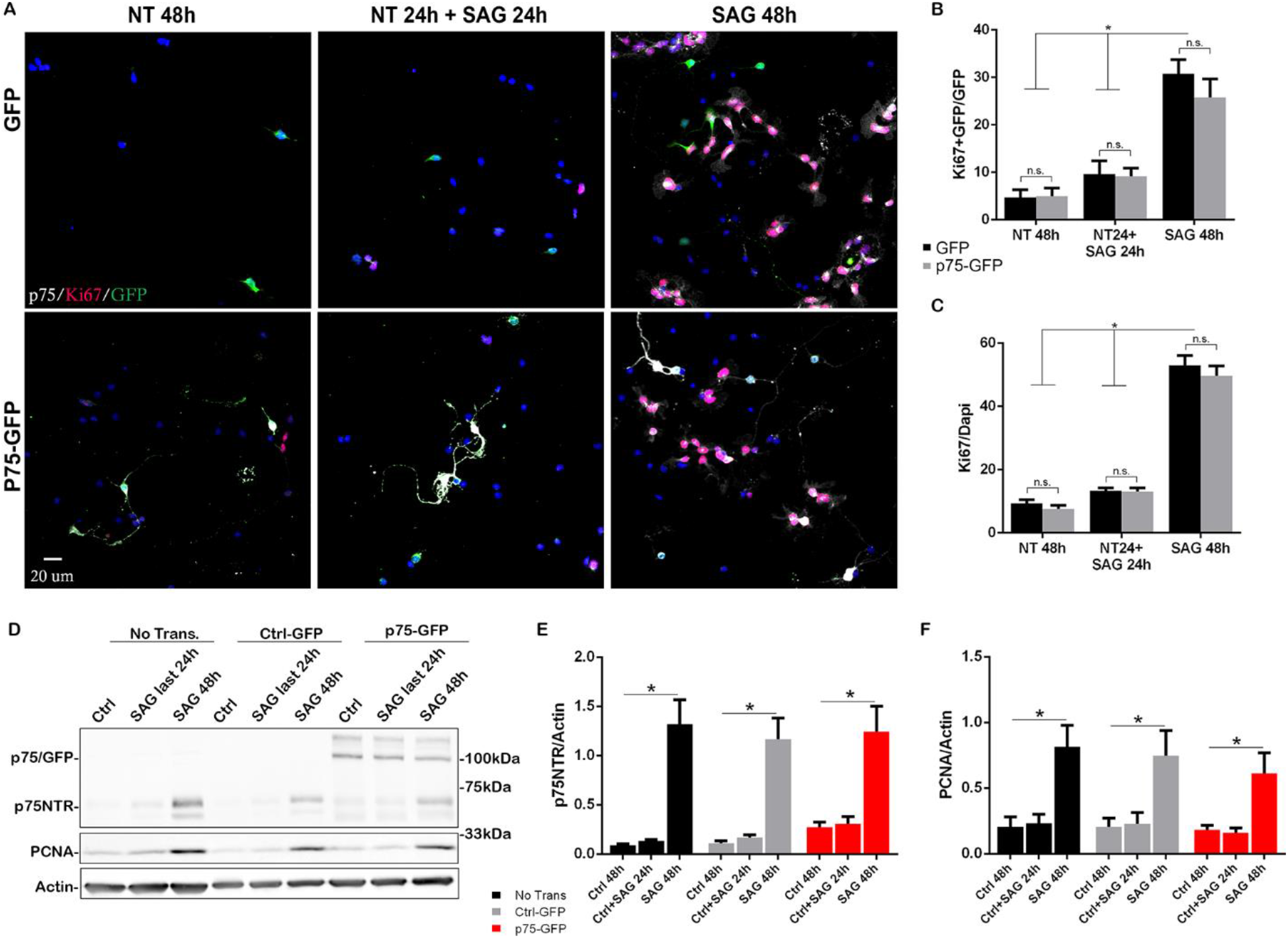
Overexpression of p75NTR is not sufficient to maintain granule cells in a proliferative state. (A) Immunostaining analysis of the expression of p75NTR (white), GFP (green), and Ki67 (magenta) in granule cell cultures transfected with Ctrl-GFP (1^st^ and 3^rd^ rows) or p75-GFP (2^nd^ and 4^th^ rows). Top two panels, transfected cells were maintained in SAG for 48h. Bottom-two panels’ cells were maintained without SAG for the first 24 h in culture, and in the presence of the mitogen only in the last 24h in culture. (B) Quantification of the transfected cells that are proliferating, expressed as the double label Ki67+GFP over the total number of transfected cells (GFP + cells). Two-way ANOVA, N = 3, error bars indicate SEM. (C) Quantification of the total number of proliferating cells, expressed as the percentage of Ki67+ cells over the total number of cells (Dapi). Two-way ANOVA, N = 3, error bars indicate SEM. (D) Western blot analysis of the expression levels of p75NTR and PCNA in granule cell cultures, first three lanes no transfected cells, last six lanes’ cells transfected with Ctrl-GFP or p75-GFP. (E and F) Quantification of the expression levels of p75NTR or PCNA relative to actin. Two-way ANOVA, N = 4, error bars indicate SEM.

### P75NTR prevents CGN migration in vitro

The lack of proliferation in response to SAG observed in cells transfected with p75NTR, suggests that the receptor is not involved in maintaining the cells in a proliferative state, ready to respond to a mitogen. Considering the sharp downregulation of p75NTR observed in migratory cells (Fig 1 and 2), an alternative explanation would be that the presence of p75NTR in proliferating GCPs prevents the onset of migration of this cell population. To assess whether the expression of p75NTR prevents granule cell migration, we used a trans-well migration assay and compared the intrinsic migratory activity of dissociated granule cells obtained from P7 WT and p75NTR-/- rat pups. To evaluate the intrinsic migratory properties of each of the genotypes, no ligand was added to the trans-well compartments. The p75NTR-/- cells showed a 2-fold increase in the number of migrating cells compared to WT cells (Fig 5A-B).

**Figure 5:**
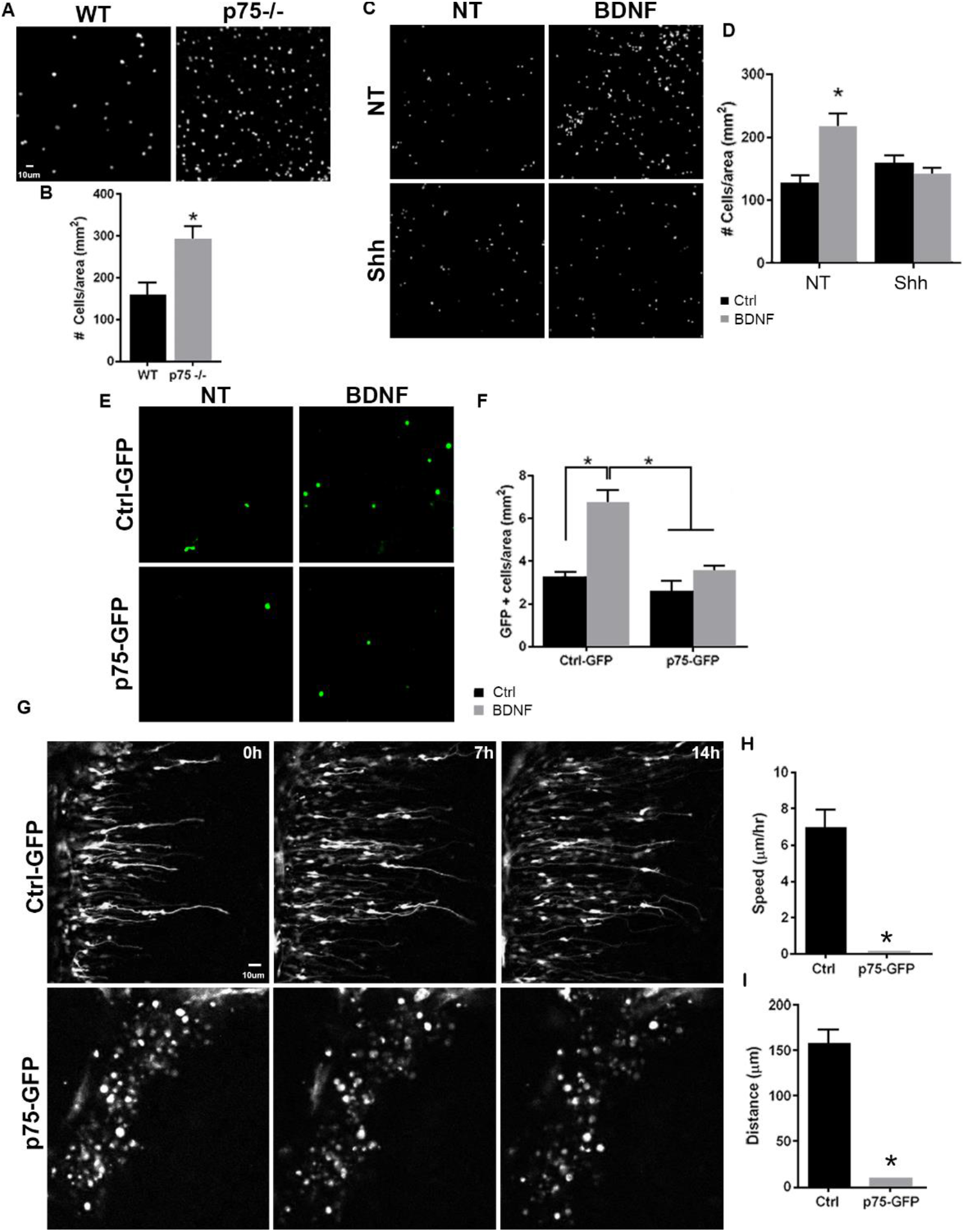
p75NTR prevents migration of the granule cells. (A) Migration analysis using transwell assay. Dapi immunostaining of cells obtained from WT (top) or p75NTR-/- (bottom) P7 rat pups. (B) Quantification of CGN migration express as the density of Dapi + cells at the bottom of the filter after 24 h. No ligand was added to stimulate the migration. One-way ANOVA, N = 4, error bars indicate SEM. (C) Migration analysis using transwell assay in cells exposed to Shh (in the top and bottom compartment) and BDNF (bottom compartment). (D) Quantification of CGN migration expressed as the density of Dapi + cells at the bottom of the filter after 24 h. Two-way ANOVA, N = 4, error bars indicate SEM. (E) Migration analysis using transwell assay in cells transfected with Ctrl-GFP or p75-GFP construct and expose to BDNF in the bottom compartment. GFP immunostaining of the transfected cells. (F) Quantification of CGN migration expressed as the density of transfected GFP+ cells at the bottom of the filter after 24 h of BDNF exposure. Two-way ANOVA, N = 4, error bars indicate SEM. (G) Time-lapse pictures from cerebellar organotypic slices were obtained from P7 rat pups and transfected with Ctrl-GFP (Top panel) or p75NTR-GFP (bottom panel). (H) Mean migration speed is expressed as the total distance migrated over the total time of the experiment. Unpaired t test, N = 3, error bars indicate SEM. (I) Total distance migrated is expressed as the mean distance migrated per cell. Unpaired t test, N = 3, error bars indicate SEM.

The previous data demonstrated that mitotic GCPs maintain high levels of p75NTR, while postmitotic cells expressing neuronal differentiation markers such as DCX or βIII tubulin completely downregulate p75NTR expression (Fig 2 and 3), supporting our hypothesis that p75NTR may serve to retain the proliferating progenitors in the mitogenic environment of the EGL. To test the possibility, using the trans-well assay we exposed the cells to Shh (5C bottom panels, 5D) to maintain them in a proliferative state expressing high levels of p75NTR, and stimulated migration using BDNF (5C right panels, 5D), a well-known chemoattractant for the postmitotic cerebellar granule neurons (CGN, Borghesani et al., 2002; Zhou et al., 2007). In the absence of Shh, BDNF stimulated the migration of CGN in agreement with the previously mentioned reports (Fig 5C-D), whereas, in the presence of Shh, which promotes proliferation and maintains elevated levels of p75NTR, BDNF did not induce migration (Fig 5C-D), indicating that only the post-mitotic CGNs that have down-regulated p75NTR responded to BDNF with increased migration. An alternative explanation for these findings could be that Shh maintains cells in a proliferative state, and restricts CGN migration independent of p75NTR. To test this possibility, we transfected CGNs to overexpress p75NTR and exposed the cells to BDNF in a trans-well assay. Our results showed an increase in migration in response to BDNF in the CGN transfected with a control GFP construct (Fig 5E top panel, 5F), similar to the CGN response to BDNF in WT non-transfected cells (Fig 5C-D). However, the chemotactic effect of BDNF was lost in the CGN overexpressing p75NTR (Fig 5E bottom panel, 5F), suggesting that p75NTR is sufficient to prevent CGN migration even in the presence of a chemotactic signal such as BDNF.

### P75NTR expression prevents CGN migration in cerebellar slices

To directly evaluate the effects of p75NTR expression during CGN migration in cerebellar tissue, we transfected p75NTR into the EGL in cerebellar organotypic slices and tracked migrating CGNs using timelapse microscopy. P7 rat cerebellum slices were electroporated either with a ctrl-GFP or p75NTR-GFP construct, specifically into the EGL. In slices transfected with the Ctrl-GFP vector, we observed cells migrating towards the IGL (Fig 5G top panels, video 1). The migration parameters, speed, and distance, were consistent with previously reported values (Fig 5H-I) (Yacubova & Komuro, 2002). These cells displayed the classic morphology of migrating neurons, a leading process extended towards the IGL was observed and the soma showed a “saltatory” migration pattern, i.e., alternating stationary and displacement periods of the cell body (Yacubova & Komuro, 2002). However, when p75NTR was overexpressed, the transfected neurons remained in the EGL. In contrast to the control conditions, these cells maintained a rounded morphology and never extended a process in any direction, resembling a nonmigrating neuron (Fig 5G bottom panels, video 2). These results strongly support the possibility that the presence of p75NTR is sufficient to prevent granule cell migration, and the receptor needs to be downregulated before the cells start to migrate.

### The absence of p75NTR in GCPs increases granule cell migration in vivo

During cerebellar development, p75NTR is expressed in GCPs and Purkinje cells (see Fig 1). To analyze the specific role of p75NTR in the transition of GCPs from a proliferating to a migrating population *in vivo*, we mated p75NTR^fl/fl^ mice (Bogenmann et al., 2011) with Atoh1^Cre^ mice (Jackson Lab). In the cerebellum, these mice lack p75NTR expression specifically in the EGL while maintaining WT expression of the receptor in the Purkinje cells (Zanin et al., 2016). We injected EdU into P7 mice and waited for 24 or 48 h before perfusing the animals to allow the dividing cells in the EGL to incorporate EdU and eventually exit the cell cycle and migrate toward the IGL. Cells positive for EdU, located in the IGL, are postmitotic GCPs that migrated in the time between the EdU injection and euthanizing the animal. At both time points, 24 and 48 h, a significantly increased number of EdU+ cells were found in the IGL of the p75^fl/fl^; Atoh1^Cre^ mice compared to WT animals (Fig 6A-C). Developmental differences between anterior and posterior lobes have been previously reported (Martinez et al., 2013), however, we observed a significant increase of EdU+ cells in the anterior lobes (lobule 5) and the posterior lobes (lobules 6 and 9) of p75^fl/fl^; Atoh1^Cre^ mice, suggesting a general mechanism of p75NTR inhibition of migration in the entire cerebellum (Fig 6B-C). PAX6 staining confirmed the cerebellar granule neuron identity of the EdU+ cells in the IGL (Fig. 6D). Cells positive for EdU in the white matter were negative for PAX6, therefore excluded from the analysis (Fig 6D dotted line).

**Figure 6:**
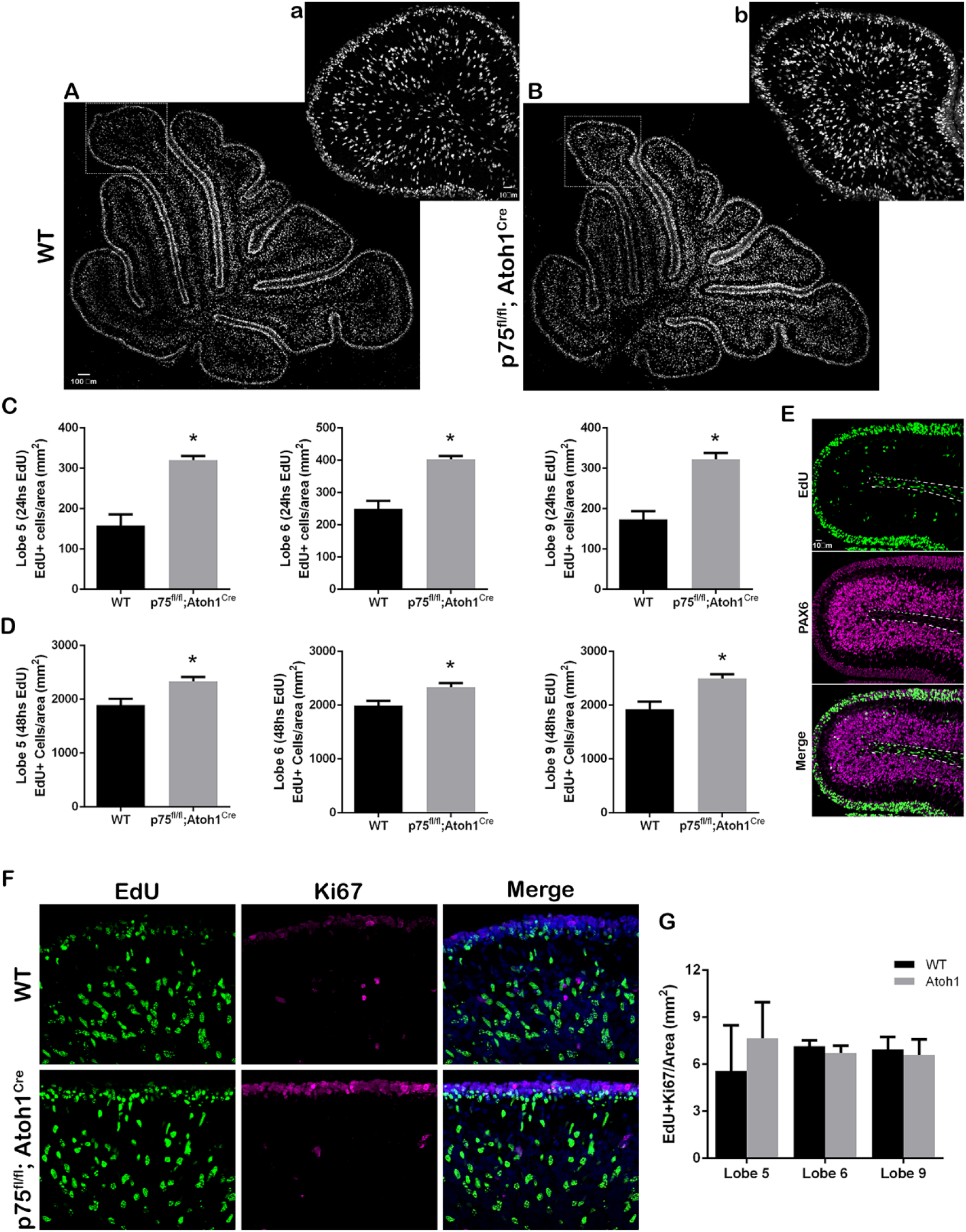
p75NTR prevents CGN migration *in vivo.* (A-B) Immunohistochemistry of cerebellar sections from (C) P8 and (D) P9 mouse pups injected with EdU 48h before euthanasian the animal. (a-b) High magnification of the inset shown in left panels. (C and D) quantification of the density of migrating cells expressed as the total number of EdU positive cells in the IGL per mm^2^. (C) P8 mouse pups injected with EdU 24h before euthanasian the animal. Unpaired t test, WT N = 6, p75^fl/fl^; Atoh1^Cre^ N = 4, error bars indicate SEM. (D) P9 mouse pups injected with EdU 48h before euthanasian the animal. Unpaired t test, WT N = 9, p75^fl/fl^; Atoh1^Cre^ N = 9, error bars indicate SEM. (E) Cerebellar section from a P8 mouse injected with EdU 24h before euthanasian the animal. EdU in green, Pax 6, granule cell marker, in magenta. (F) Immunohistochemistry of cerebellar sections from P9 mouse pups injected with EdU 48h before euthanasian the animal. EdU in green, Ki67 (proliferation marker) in magenta. (G) Quantification of the migrating cells that continued to express Ki67, express as the density of EdU/Ki67 double-label cells in the IGL. One-way ANOVA, N = 3, error bars indicate SEM.

The increased presence of granule cells in the IGL observed in the p75^fl/fl^; Atoh1^Cre^ mice might be due to abnormal early migration of the GCPs if the cells started the migratory process before they exited the cell cycle. To assess whether cells migrated inappropriately while still in the cell cycle, we stained the EdU injected animals with Ki67, a proliferation marker present in actively proliferating cells during the entire cell cycle (Gerdes et al., 1983; Scholzen & Gerdes, 2000). A double-labeled cell (EdU/Ki67 positive cell) in the IGL would indicate premature GCP migration while the cells were still proliferating. Our results showed that in WT as well as in p75^fl/fl^; Atoh1^Cre^ animals, the number of EdU/Ki67 positive cells present in the IGL was close to zero, and no difference was observed between the genotypes (Fig 6F-G), indicating that no premature cell migration of proliferating GCPs was observed when we removed p75NTR, therefore cells still exited the cell cycle before migration.

### RhoA inhibits GCP radial migration

RhoA is a member of the Rho GTPase family of proteins which play an important role in cell migration via cytoskeletal regulation (Ridley, 1995; Ridley & Hall, 1992). RhoA activity is known to be regulated by p75NTR, and granule cells from p75NTR-/- mice show reduced levels of active RhoA (Fig 7A). Therefore, to test the involvement of RhoA in CGN migration, we used the Rho kinase inhibitor, Y27632, to block the RhoA signaling pathway and analyzed CGN migration using the trans-well assay. Even in the absence of a chemoattractant, inhibition of the RhoA pathway significantly increased the number of migrating cells compared to the control condition (Fig 7B, black bars), similar to the p75NTR knockout CGNs. Moreover, the inhibition of Rho-kinase did not affect the ability of BDNF to promote migration, suggesting that RhoA is not required for CGN migration (Fig 7B gray bars). Since inhibition of RhoA signaling promoted migration, we assessed whether RhoA must be downregulated for CGN migration to occur. Organotypic cultures transfected with Ctrl-GFP migrated normally when exposed to a Rho kinase inhibitor (Fig 7C, Video 3), confirming that RhoA activity is not necessary for migration to occur (compare Video 1 and 3). However, overexpression of a constitutively active form of RhoA (CA-RhoA) in the EGL of P7 cerebellar slices inhibited migration toward the IGL (Fig 7D, video 4), similar to the overexpression of p75NTR-GFP (see Fig 5E), and the majority of the cells maintained a rounded shape with no extended process in the transfected cells (Fig 7D, video 4). After finishing the time-lapse video, the cerebellar slices were fixed and stained for DCX, to identify the migrating cells, and GFP to identify the transfected cells overexpressing the CA-RhoA (Fig 7E). The cells expressing the CA-RhoA remained in the EGL, and no GFP was detected in the IGL. Consistent with the anti-migratory role of RhoA, no colocalization between the CA-RhoA cells and DCX was observed (Fig 7E).

**Figure 7:**
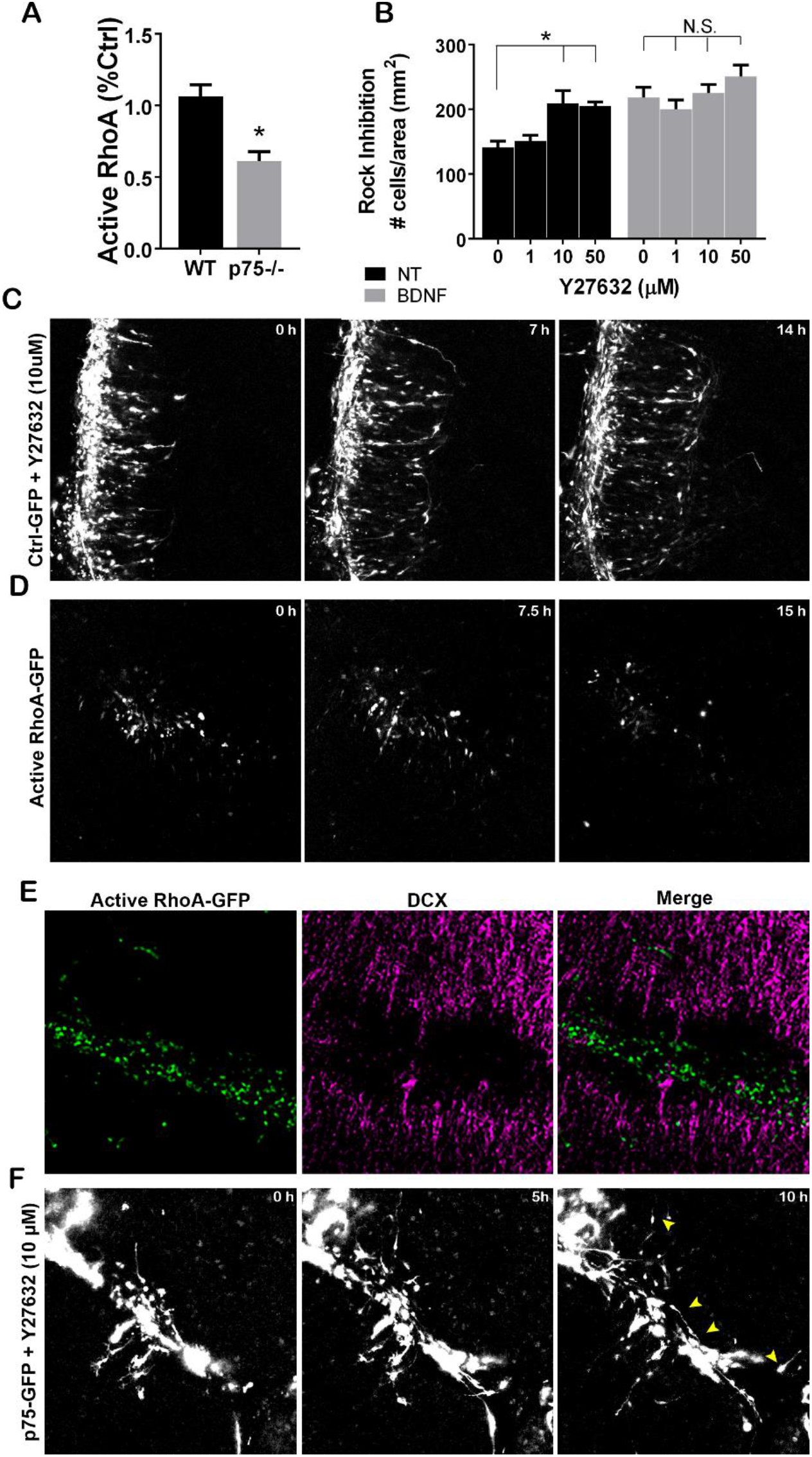
(A) Quantification of the levels of active RhoA in granule cell cultures from WT or p75-/- animals after 48h. Unpaired t Test, N = 4, error bars indicate SEM. (B) Migration analysis using transwell assay in cells exposed to the Rock inhibitor, Y27632 (in the top and bottom compartment) and BDNF (bottom compartment). Two-way ANOVA, N = 5, error bars indicate SEM. (C) Timelapse pictures from cerebellar organotypic slices from P7 rat pups, transfected with Ctrl-GFP in the presence of the RhoA inhibitor, Y27632. (D) Time-lapse pictures from cerebellar organotypic slices from P7 rat pups, transfected with a RhoA constitutive construct. (E) Immunostaining of the organotypic slice shown in D. RhoA-GFP (green), DCX (magenta). (F) Time-lapse pictures from cerebellar organotypic slices from P7 rat pups, transfected with a p75NTR-GFP construct and expose to Rock Kinase inhibitor (Y27632). Arrowheads indicate the migrating neurons where a leading process is observed.

### P75NTR inhibits GCP migration via RhoA Activation

p75NTR can interact with Rho-GDI at the plasma membrane of GCPs, helping to maintain high levels of active RhoA in a ligand-independent fashion (Yamashita et al., 1999; Yamashita & Tohyama, 2003). We have previously demonstrated that active RhoA was necessary for the GCPs to remain in the cell cycle, and inhibition of RhoA elicited cell cycle exit of GCPs, which was correlated with a downregulation of p75NTR (Zanin et al., 2019). Furthermore, p75-/- GCPs showed a significant reduction in the levels of active RhoA compared to WT cells (Fig 7A). These results suggest that the elevated levels of p75NTR present in the proliferating GCPs might prevent cell migration via RhoA activation, and retain the GCPs in the mitogenic environment of the EGL. To address this possibility, we transfected cerebellar slices with a p75-GFP construct and added the RhoA inhibitor Y27632 directly into the bath. Our results showed that inhibiting RhoA was sufficient to induce migration of the CGN (Fig7F, and video 5). These results confirm that the inhibition of CGN migration induced by p75NTR requires the activation of RhoA.

### Cdc42 is necessary for GCP radial migration

In postmitotic GCPs, Cdc42, another member of the Rho GTPase protein family, is required to extend the leading process that will guide the cells toward the IGL (Govek et al., 2018). To confirm that Cdc42 is required for GCP migration, we blocked Cdc42 activity using ML141 and quantified GCP migration using a trans-well assay. In the control condition (without BDNF as a chemoattractant), there was a significant reduction in GCP basal migration (Fig S2 black bars). Even in the presence of BDNF, we observed a dosedependent reduction in GCP migration after inhibiting Cdc42 (Fig S2 gray bars) confirming that Cdc42 is required for GCP migration.

Using cerebellar slices transfected with the GFP-ctrl construct, we evaluated the migration of GCPs after the addition of the CdC42 inhibitor to the culture media. After blocking Cdc42, the majority of GFP+ cells remained in the EGL and almost no radial migration towards the IGL was observed (Fig S2B, video 5), similar to the results observed after overexpressing p75NTR-GFP (see Fig 5E). These results indicate the requirement for Cdc42 in GCP migration, confirming previous studies (Govek et al., 2018). It is worth noting that tangentially migrating neurons can still be observed after blocking CdC42 (video 5, top right corner) suggesting a different mechanism between these two types of neuronal migration.

## Discussion

Neuronal migration is one of the fundamental mechanisms by which the nervous system acquires its final shape and function. Defects in neuronal migration often result in neuronal cell death or misplacement of a neuronal population, leading to a variety of disorders including epilepsy, mental retardation, and cortical dysplasia, among other neurological disorders (Copp & Harding, 1999; Moffat et al., 2015; Pilz et al., 2002). Although there is considerable emphasis on understanding the mechanisms that promote neuronal migration, as well as identifying potential signals that support cell guidance, little is known about the mechanisms that restrict migration and regulate the timing of the onset of migration. The balance between promoting and preventing migration is critical for the timely coordination of the developing nervous system since it positions the correct neuronal types at a precise time for the establishment of functional neuronal connections. In the cerebellum, it has been proposed that the external granule layer (EGL) represents a mitogenic niche permissive for granule cell proliferation, and migration away from this niche results in granule cell differentiation (Choi et al., 2005). Consistent with this hypothesis, in the present work we identify a novel function for p75NTR in restricting the initiation of migration of cerebellar granule cell precursors (GCP), providing a mechanism that explains how GCPs are retained in the EGL.

### P75NTR does not maintain GCPs in a proliferative state

Shh is secreted by Purkinje cells and is a well-established mitogen for GCPs in the EGL during early cerebellar development. p75NTR is expressed in all the proliferating GCPs in the external EGL and is downregulated in postmitotic cerebellar granule neurons (CGN). The same expression pattern is observed *in vitro*, where in the presence of Shh the cells continued to proliferate and express p75NTR. Moreover, stimulation of GCP cell cycle exit using proNT-3 or PACAP, induced a reduction in p75NTR expression together with the cessation of proliferation. ProNT-3 and PACAP both induced cell cycle exit even in the presence of Shh, suggesting that GCP cell cycle exit is an active mechanism depending on the proper signals and not just the absence of the mitogen. Since p75NTR is highly expressed in all the proliferating GCPs, one possibility was that the expression of this receptor maintains the cells in a permissive state for proliferation. In this study, we confirmed that continued expression of p75NTR was not sufficient to maintain the GCPs in a proliferative state, once the mitogen was absent for at least 24h and the cells had begun to differentiate since re-addition of Shh did not evoke cell cycle re-entry in cells that continued to express p75NTR.

### P75NTR prevents granule cell migration

The well-defined boundary between the migrating granule cells expressing DCX and the proliferating GCPs expressing p75NTR suggested that the receptor might be involved in regulating the onset of GCP migration. In this study, we demonstrated that the presence of p75NTR was sufficient to prevent the migration of GCPs. Using a trans-well assay we observed that p75-/- cells have an intrinsically higher rate of migration compared to WT cells. Moreover, cells that were grown in the presence of Shh, which induces proliferation of the GCPs while maintaining elevated expression levels of p75NTR, were sufficient to reduce the migration of GCPs, even in the presence of BDNF (a known chemotactic signal for these cells). Although maintaining the cells in the cell cycle may prevent the onset of migration by a variety of mechanisms, we demonstrated that the overexpression of p75NTR in CGNs was sufficient to block migration even in the presence of BDNF. These observations raise the possibility that the high expression of p75NTR in the presence of Shh serves to keep the cells in the proliferative environment of the EGL.

To further test this possibility, we used time-lapse microscopy in cerebellar organotypic cultures, maintaining the cells in the proper tissue environment and in the presence of the molecular signals that regulate GCP proliferation and migration. Overexpression of p75NTR was sufficient to block GCP migration from the EGL. Interestingly, these cells maintained a rounded shape and no process was observed in any direction, unlike the cells in the control conditions where a clear leading process was observed in the cell edge facing the internal granule layer. Since elaboration of the leading process is critical for the onset of GCP migration, the continued expression of p75NTR may prevent the changes in cytoskeletal dynamics required for migration to occur.

During cerebellar development, p75NTR is expressed in the GCPs throughout the entire EGL surface, including every lobe of the vermis and the hemispheres (Balaei et al., 2016; Carter et al., 2003; Zanin et al., 2016). Our *in vivo* experiments demonstrated that the absence of p75NTR was sufficient to increase GCP migration in both the anterior and posterior lobes, suggesting a common mechanism regulating the onset of migration throughout the entire cerebellum. These findings are particularly interesting since different areas and lobes of the cerebellum are associated with different behaviors (Buckner, 2013; Buckner et al., 2011; Voogd & Glickstein, 1998). Since the absence of the receptor affected the entire cerebellum, our findings reveal an important role for p75NTR, dysregulated migration in the absence of the receptor can potentially impact a wide range of behaviors.

### Rho A is required to prevent GCP migration

RhoA has been associated both positively and negatively with cell migration via cytoskeletal regulation, this mechanism has been demonstrated in different cell populations including cerebellar granule cells (Amano et al., 2010; Bito et al., 2000; Fukata et al., 2003). In CGNs, p75NTR can constitutively maintain elevated levels of active RhoA in the absence of ligand (Yamashita et al., 1999; Yamashita & Tohyama, 2003). In our previous study, we demonstrated that proNT-3 induced cell cycle exit of the CGPs (Zanin et al., 2016) and at the same time it induced a reduction in the levels of active RhoA (Zanin et al., 2019). Consistent with these results, we showed that p75NTR-/- cells have a reduced level of active RhoA compared to WT cells, and inhibition of RhoA was sufficient to promote the migration of GCPs. Moreover, overexpressing a constitutively active form of RhoA prevented the migration of GCPs, while blocking RhoA activity in organotypic cultures did not affect GCP migration. Taking into consideration our previous and current findings, the inhibition of migration induced by p75NTR might be due to the maintenance of active RhoA observed in the presence of the receptor. Upon cell differentiation, the levels of p75NTR are reduced inducing a reduction in the activity of RhoA, allowing the cells to migrate.

Although p75NTR can promote migration of multiple cell populations including neural crest cells (NCC, Zanin et al., 2013), neuronal progenitors in the rostral migratory stream (Snapyan et al., 2009) or even certain types of cancers (Wislet et al., 2018), our findings demonstrated an anti-migratory effect for this receptor in granule cell precursors. A previous study proposed that proBDNF negatively regulates the migration of granule cell precursors and this effect is mediated by p75NTR. In that work, the author described that p75NTR-/- mice have reduced CGNs migration from the EGL. Although we do not discard the involvement of a neurotrophin ligand to regulate migration, our results demonstrated that an un-liganded p75NTR is responsible for preventing CGNs migration, while removing the receptor is sufficient to stimulate the migration of the CGNs. These apparently contradictory results might be explained due to the wide range of signaling pathways that can be regulated by p75NTR. Depending on the receptor complex, p75NTR can be activated by mature neurotrophins or proneurotrophins, it can also modulate the affinity of mature neurotrophins for the Trk family of receptors or it can modulate the activation of intracellular signals in a ligand-independent manner. All these possibilities highlight the complexity of p75NTR signaling and the urgent necessity to address them in a cell-specific context, as they could provide an understanding of normal embryonic development and adult brain function while providing new targets for potential novel therapeutics.

## Acknowledgments

The p75-GFP construct was generously provided by Moses Chao (NYU Medical School). This work was funded by NIH/NINDS 1R56NS094589 to WJF, and Rutgers Busch Biomedical Grant AWD00009650 to JPZ.

## Author contributions

J.P.Z. and W.J.F. designed the study, J.P.Z. performed the experiments, J.P.Z. and W.J.F. wrote the manuscript.

## Competing Interests

The authors declare no competing interests.

## Materials and methods

### Primary cerebellum cell cultures

All animal studies were conducted using the National Institutes of Health guidelines for the ethical treatment of animals with the approval of the Rutgers Animal Care and Facilities Committee. Cerebella were removed under sterile conditions from P7 pups after euthanizing with CO2. Meninges and small blood vessels were removed under a dissecting microscope. The tissue was minced and dissociated using the papain dissociation kit (Worthington LK003150). Dissociated neurons were plated onto 24 well plates (1 × 10^5^ cells in 300 μl of serum-free media) or 6 well plates (1.5 × 10^6^ cells per well in 1 ml of serum-free media) coated with poly-D-lysine (0.1mg/ml). Serum-free medium consisted of 1:1 MEM and F12, with glucose (6mg/ml), insulin (2.5mg/ml), putrescine (60 μM), progesterone (20nM), transferrin (100μg/ml), selenium (30nM), penicillin (0.5U/ml) and streptomycin (0.5μg/ml).

### Immunohistochemistry

Animals were deeply anesthetized with ketamine/xylazine and perfused with 4% PFA/PBS. Brains were removed and postfixed in 4% PFA/PBS overnight at 4°C, then cryopreserved with 30% sucrose. Sections (12μm) were cut using a Leica cryostat and mounted onto charged slides. Sections were permeabilized with 0.5% Triton in PBS for 20 min and blocked with 1% BSA and 5% donkey serum in PBS for 1h at room temperature. EdU stained was developed following manufacturer instruction (Thermo Fisher Scientific Cat# C10337). Antibody staining was started immediately after finishing EdU staining. Primary and secondary antibodies were prepared in 1% BSA. Sections were incubated with primary antibodies overnight at 4°C in a humidified chamber. Antibodies used were: Ki67 (Abcam 15580, RRID: AB_443209, 1:500), anti-p75 (R&D AF367, RRID: AB_2152638, 1:500), anti-p75 (Millipore MAB365, RRID: AB_2152788, 1:1000) anti-PAX-6 (BD Bioscience, RRID: AB_10715442, 1/500), anti-DCX (Abcam, RRID: AB_732011, 1/500) and anti-TAG1 (Cell Signaling, RRID: AB_2798922, 1/500). All secondary antibodies were diluted 1:1000 and incubated for 1h at RT. Sections were mounted using DAPI Fluoromount-G (Southern Biotech #0100-20). Controls for immunostaining included incubation with secondary antibodies in the absence of primary antibodies.

### Immunocytochemistry

Cells were fixed in 4% PFA/PBS for 20 min at room temperature and permeabilized with 0.5% Triton in PBS for 15 min and blocked with 1% BSA and 5% donkey serum in PBS for 1h at room temperature. EdU stained was developed following manufacturer instruction (Thermo Fisher Scientific Cat# C10337). Antibody staining was started immediately after finishing EdU staining. Primary and secondary antibodies were prepared in 1% BSA. Cells were incubated with primary antibodies overnight at 4°C in a humidified chamber. Antibodies used were: anti-p75 (R&D AF367, RRID: AB_2152638, 1:500), anti-DCX (Abcam, RRID: AB_732011, 1/500), Ki67 (Abcam 15580, RRID: AB_443209, 1:500), TUJ1 (Promega, RRID: AB_430874, 1/1000), anti-GFP (Sigma, RRID: AB_259941, 1/500). All secondary antibodies were diluted 1:1000 and incubated for 1h at RT. Sections were mounted using DAPI Fluoromount-G (Southern Biotech #0100-20). Controls for immunostaining included incubation with secondary antibodies in the absence of primary antibodies.

### Cell culture transfection

P7 dissociated granule cells were obtained as described above and transfected using Nucleofector II (Lonza) following manufacturer specifications. Briefly, 2 to 4 × 10^6^ cells were spun down at 80 g for 10 min. The cell pellet was resuspended in 100 μl of transfection buffer (Lonza). The corresponding DNA construct to be transfected was added and mixed in the resuspended cells at a concentration of 1μg DNA/million cells. Immediately after mixing, the cells were electroporated using a Nucleofector II (Lonza), using the G-013 program. After electroporation, 500 μl of SFM media supplemented with 1% B27 was added to the cells and transfered to a 15 ml conical tube. The cell suspension was left to recover for 5 min in the incubator (37°C, 5% CO2). The cells were plated at a concentration of 1 × 10^5^ per well in a 24 well plate with Poly D lysine coated coverslip.

### Organotypic culture and cell migration assay

After removing and washing the cerebellum, DNA constructs were injected between lobes and electroporated using the following parameters: 5 pulses, 30V each, 50 ms ON-950 ms OFF (Harvard apparatus ECM 830 BTX). After electroporation, the cerebellum was sliced into 280 μm sections using a tissue chopper machine (McIlwain Tissue Chopper 800 series Vibratome). The sliced cerebellum was placed in a 10 cm Petri dish with 6 ml of media and gently rocked to fully separate all the sections. 3 to 4 slices were transferred to a filter insert (Millipore, Millicell cat# PICM0RG50) with 1 ml SFM media in the bottom well. Sections were incubated O.N. at 37°C and 5% CO2. The next day the insert was transferred to a 35mm glass-bottom dish (Mat Tek Cat# P35G-0-10-C) for live image recording for 15 h, using a Zeiss 510 META microscope.

### Transfection constructs

The RhoA constitutively active constructs were purchased from Addgene (pcDNA3-EGFP-RhoA-Q63L; Plasmid #12968). The ctrl-GFP construct was purchased from Lonza (pmaxGFP™ Vector). P75NTR-GFP construct was a kind gift from Dr. Moses Chao.

### pGEX-2T-RhoA-T19N (Plasmid #12960)

#### Trans-well migration assay

Trans-well assay was performed following manufacturer guidance (Costar Cat # 3422). Briefly, granule cells were plated in the top of the trans-well insert 8 μm pore size (Costar Cat# 3422), at a final density of 5 × 10^5^ cells/insert. The final volume for the top chamber was 100 μl and 600 μl for the bottom chamber. For control treatments, no ligand was added to the bottom chamber, for chemotactic analysis, BDNF was added to the bottom chamber to a final concentration of 50 ng/ml. The insert was incubated for 24hs at 37°C and 5% CO2. Cells were fixed using 4% ice-cold paraformaldehyde for 20 min and washed 3 times with PBS. Using a Q-tip, the cells on the top of the filter were removed by scraping the surface. Cells at the bottom of the filter were stained using 1μg/ml of Dapi for 3 min at R.T. Almost? 90% of the filter surface was imaged using a Nikon E1000 with a 4X magnification. The total number of cells migrated was quantified using ImageJ V1.51.

For the inhibition experiments, the specific inhibitor was added to the bottom and top compartment. All the inhibitors were added from the time of plating the cells and maintained for 24 h. For RhoA inhibition, we use Rock inhibitor Y27632 (Calbiochem Cat# 688000) to a final concentration of 1, 10, and 50μM. For CdC42 inhibition, we use ML141 (Sigma Cat# 217708) to a final concentration of 1, 2.5, 5, and 10 μM.

#### In vivo migration assay

P7 mouse pups were injected with 50 mg/kg EdU intraperitoneally. After 24 or 48 h, animals were deeply anesthetized with ketamine/xylazine and perfused with 4% PFA/PBS. Brains were removed and postfixed in 4% PFA/PBS overnight at 4°C, then cryopreserved with 30% sucrose. Sections (12 μm) were cut using a Leica cryostat and mounted onto charged slides. EdU stained was developed following manufacturer instruction (Thermo Fisher Scientific Cat# C10337). Stained for PAX6 was started immediately after finishing EdU staining, as described above. Sliced sections were imaged using a Nikon E1000 with a 10X magnification. The total number of EdU+ and PAX6+ cells was quantified using ImageJ V1.51.

#### GTPase activation analysis

Cells were obtained from P7 WT and p75-/- pups and cultured as described above for 48h. Cells were processed for G-Lisa analysis according to the manufacturer’s instructions. G-Lisa for RhoA (Cytoskeleton Inc., #BK124).

#### Western Blot

Cultured cells were washed with ice-cooled PBS and homogenized using 1% NP40, 1% Triton, and 10% glycerol in TBS buffer (50mM Tris, pH 7.6, 150 mM NaCl) with a protease inhibitor cocktail (Roche Products, 11 836 153 001). Proteins were quantified using the Bradford assay (Bio-Rad 500-0006) and equal amounts of protein were run on SDS gels and transferred to a nitrocellulose membrane. To ensure equal protein levels, blots were stained with Ponceau before incubation with antibodies. The blots were then rinsed and blocked in 5% nonfat dried skim milk in TBS-T for 1 h at RT. Blots were incubated with primary antibodies diluted 1:1000 in 1% BSA in TBS buffer overnight at 4°C. The blots were washed with TBS-T 3 x 10 min each and incubated with Licor secondary antibody for 1 h at RT. All secondary antibodies were diluted at 1:10,000. Membranes were washed 3 x 10 min each in TBS-T. The membranes were analyzed using Licor Odyssey infrared imaging system (LICOR Bioscience). To confirm equal protein levels, blots were reprobed for actin. All analyses were performed at least three times in independent experiments. Bands were quantified using Image J Version 1.51s.

#### Tissue clarification

The clarification of cerebellar slices was done following the iDISCO protocol (version 2016). Briefly, after the time-lapse experiment was finished, the membrane filter that contained the slice was cut using a scalpel blade. The section was fixed at room temperature (RT) for 1h and washed with PBS RT 3 times x 20 min. The section was permeabilized with 1xPBS/0.5% TritonX-100/ 20%DMSO at RT for 2h and blocked at RT using Donkey Serum 5% and BSA 1% in PBS for 2 h. The primary antibody was incubated overnight at 4°C in a rocking station. The next day, the primary antibody was washed with PBS 3 times x 30 min. The secondary antibody was incubated overnight at 4°C in a rocking station.

For the clearing procedure sample was dehydrated in ascending methanol series:20%, 40%, 60%, 80%, 100%, 100%; for 1h each at RT. The sample was incubated in 66% dichloromethane (DCM, Sigma 27099712 × 100ml) /33% Methanol at RT for 3h, with shaking, and then incubated in 100% DCM for 2 × 15 minutes to wash the MeOH, with shaking. Finally, the sample was incubated in DiBenzyl Ether (DBE, Sigma 108014-1KG). To acquire the images, the sample was placed in a glass-bottom 35 mm petri dish cover with DBE.

#### Statistical Analysis

Experimental groups were compared using either Student’s t-test, one- or two-way ANOVA followed by Tukey’s posthoc analysis, as appropriate, p < 0.05 was considered significant. The specific statistical analysis is indicated in each figure legend.

## Notes

### Competing Interest Statement

The authors have declared no competing interest.

