## Supplemental Figures for "p75NTR prevents the onset of cerebellar granule cell migration via RhoA activation"

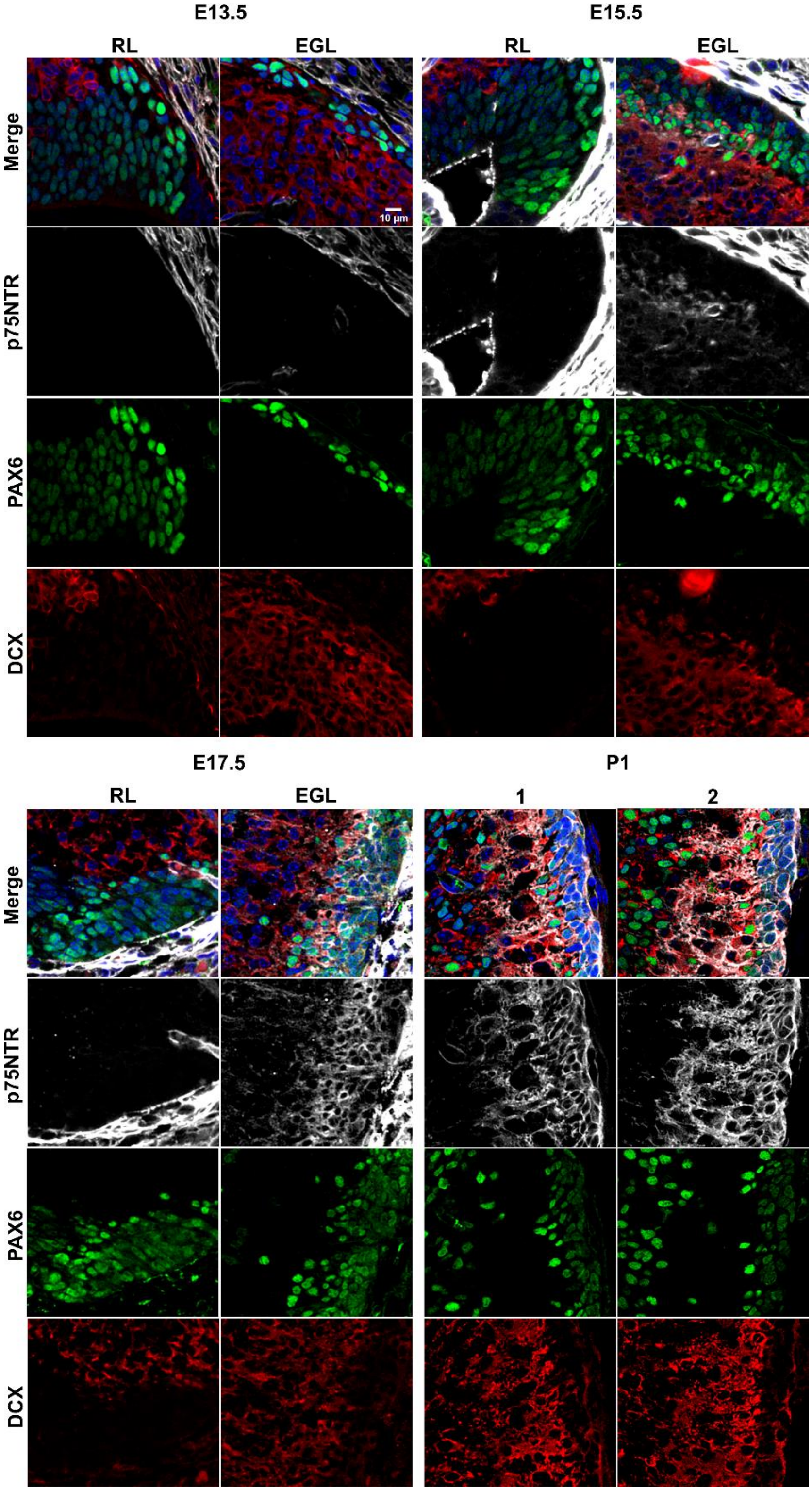


Supplemental Figure 1:
Developmental expression of p75NTR (white), Ki67 (red), and Pax6 (green) at the indicated ages. Note the high level of p75NTR in the meninges as well as the developing granule cell progenitors. RL- rhombic lip, EGL – external granule layer, M – meninges.

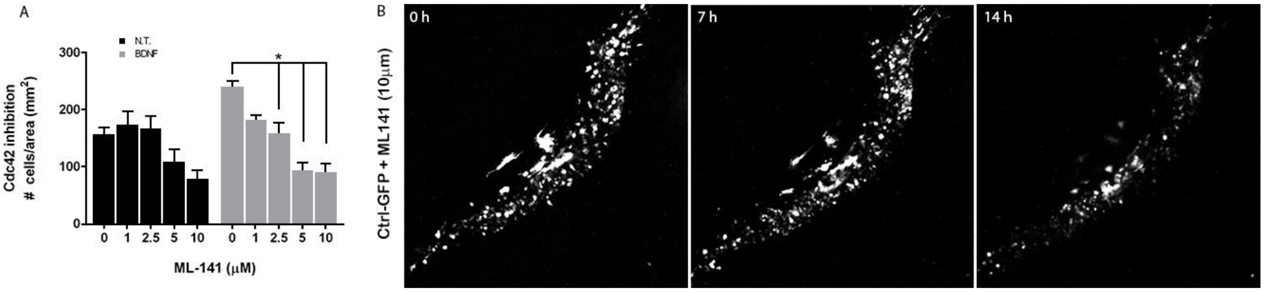


Supplemental Figure 2:

(A) Migration analysis using transwell assay in cells exposed to the CdC42 inhibitor, ML141 (in the top and bottom compartment), and BDNF (bottom compartment). (B) Time-lapse pictures from cerebellar organotypic slices from P7 rat pups, transfected with a Ctrl-GFP construct, in the presence of the CdC42 inhibitor, ML141. Two-way ANOVA, N = 3, error bars indicate SEM.
